# Retinal microvasculature and cerebral small vessel disease in the Lothian Birth Cohort 1936 and Mild Stroke Study

**DOI:** 10.1101/462507

**Authors:** Sarah McGrory, Lucia Ballerini, Fergus N. Doubal, Julie Staals, Mike Allerhand, Maria del C. Valdes-Hernandez, Xin Wang, Thomas J. MacGillivray, Alex S.F. Doney, Baljean Dhillon, John M. Starr, Mark E. Bastin, Emanuele Trucco, Ian J. Deary, Joanna M. Wardlaw

**Affiliations:** VAMPIRE project, Centre for Clinical Brain Sciences, University of Edinburgh, Edinburgh, UK; Department of Psychology, University of Edinburgh, Edinburgh, UK; Department of Neurology, Maastricht University Medical Center, Maastricht, The Netherlands; Cardiovascular Research Institute Maastricht (CARIM), Maastricht University, Maastricht, The Netherlands; Centre for Cognitive Ageing and Cognitive Epidemiology, University of Edinburgh, Edinburgh, UK; Division of Cardiovascular and Diabetes Medicine, Medical Research Institute, Ninewells Hospital and Medical School, Dundee, UK; Alzheimer Scotland Dementia Research Centre, University of Edinburgh, Edinburgh, UK; Scottish Imaging Network, a Platform for Scientific Excellence (SINAPSE) Collaboration, Edinburgh, UK; VAMPIRE project, Computing, School of Science and Engineering, University of Dundee, Dundee, UK; UK Dementia Research Institute at the University of Edinburgh, Chancellor’s Building, Edinburgh, UK

**Author notes:** These authors contributed equally (Ian J. Deary, Joanna M. Wardlaw).

## Abstract

Research has suggested that the retinal vasculature may act as a surrogate marker for diseased cerebral vessels. Retinal vascular parameters were measured using Vessel Assessment and Measurement Platform for Images of the Retina (VAMPIRE) software in two cohorts: (i) community-dwelling older subjects of the Lothian Birth Cohort 1936; and (ii) patients with recent minor ischaemic stroke of the Mild Stroke Study. Imaging markers of small vessel disease (SVD) (white matter hyperintensities [WMH] on structural MRI, visual scores and volume; perivascular spaces; lacunes and microbleeds), and vascular risk measures were assessed in both cohorts. We assessed associations between retinal and brain measurements using structural equation modelling and regression analysis. In the Lothian Birth Cohort 1936 arteriolar fractal dimension accounted for 4% of the variance in WMH load. In the Mild Stroke Study lower arteriolar fractal dimension was associated with deep WMH scores (odds ratio [OR] 0.53; 95% CI, 0.32-0.87). No other retinal measure was associated with SVD. Reduced fractal dimension, a measure of vascular complexity, is related to SVD imaging features in older people. The results provide some support for the use of the retinal vasculature in the study of brain microvascular disease.

## Introduction

Presently, the pathogenesis of cerebral small vessel disease (SVD) in humans is not completely understood, in part due to the difficulty in imaging in vivo the cerebral microvasculature. Due to the anatomical and physiological similarity of the cerebral and retinal vessels^1^, and the ease with which the retinal microvasculature can be imaged, there has been considerable interest in determining whether pathologic changes in the retinal vessels may parallel changes in the cerebral small vessels.

There is strong evidence that retinal vascular features are associated with stroke^2-3^, SVD stroke subtype^4-9^, and presence and progression of features of SVD^10-14^. However, previous analyses have focused largely on individual retinovascular signs and individual SVD markers, rather than considering retinovascular abnormalities or SVD features as a whole^11,12,15^.

We aimed to assess an array of retinal measurements in an effort to identify those most closely related to individual and total SVD burden independently of any co-association with common vascular risk factors (VRFs). We hypothesised that associations between retinovascular changes and SVD lesions would be partly explained by, but remain independent of, VRFs. We used quantitative retinal measurements, qualitative (visual ratings) and quantitative indicators of white matter hyperintensities (WMH) (WMH% volume) and SVD burden, medical-history-derived- and measured vascular risk variables, in two cohorts: the Lothian Birth Cohort (LBC1936), and the Mild Stroke Study (MSS). The LBC1936 represents a narrow age range at the ‘healthy’ end of the SVD spectrum. The MSS represents a wider age range with clinically evident brain vascular disease and a higher burden of SVD and VRFs. Analysis of these separate but complementary population samples allows us to examine the relationship between retinovascular and SVD features at different points on the SVD spectrum.

## Results

### LBC1936

In the LBC1936, 866 participants (448, 52% male) of mean age 72.5 years (SD 0.7) attended for clinical assessment; 681 participants completed brain imaging and provided WMH and SVD ratings and volumes. The 603 participants with retinal imaging measurements available for both right and left eyes provided data as the analytic sample for the present study (Table 1). Of the 603 (303, 50% male), 270 (45%) had hypertension, 58 (10%) had diabetes (Table 1), and 31% had moderate or severe periventricular WMH scores with 19% moderate or severe deep WMH scores (Table 2). Of the 603, 33 participants reported having had a stroke (of whom 11 also had evidence of stroke on imaging), and an additional 51 had imaging-only evidence of a stroke lesion, giving a total with any stroke of 84 (14%).

**Table 1.** Characteristics of the Lothian Birth Cohort 1936 and the Mild Stroke Study: M (SD) for Continuous Variables and N (%) for Categorical Variables

| Variable | Lothian Birth Cohort 1936 | Mild Stroke Study |
| --- | --- | --- |
| Sample no. | 603 | 155 |
| Age, mean (SD) | 72.5 (0.7) | 66.9 (11.4) |
| Sex |  |  |
| Female | 300 (50) | 47 (32) |
| Male | 303 (50) | 98 (68) |
| *Vascular risk factors* |  |  |
| Hypertension |  |  |
| No | 333 (55) | 58 (40) |
| Yes | 270 (45) | 87 (60) |
| Diabetes |  |  |
| No | 545 (90) | 126 (87) |
| Yes | 58 (10) | 19 (13) |
| Hypercholesterolemia |  |  |
| No | 375 (62) | 88 (62) |
| Yes | 228 (38) | 55 (38) |
| Cholesterol, mmol/L | 5.24 (1.16) | 4.99 (1.25) |
| HbA1c | 5.72 (0.60) | Blood Glucose, mmol/L<br>6.24 (3.11) |
| Smoker |  |  |
| Ex, or never | 557 (92) | 61 (42) |
| Yes | 46 (8) | 84 (58) |
| Diastolic BP, mm Hg | 78.22 (9.59) | 85.32 (13.47) |
| Systolic BP, mm Hg | 148.48 (18.99) | 150.32 (28.26) |

*Note.* BP=blood pressure; HbA1c=haemoglobin A1c; WMH=white matter hyperintensities

**Table 2.** Descriptive Statistics for Brain and retinal imaging-derived measurements in the Lothian Birth Cohort 1936 and the Mild Stroke Study: M (SD) for Continuous Variables and N (%) for Categorical Variables

| Variable | Lothian Birth Cohort 1936 | Mild Stroke Study |
| --- | --- | --- |
| *Brain measurements, mean (SD)* |  |  |
| WMH % in BTV | 1.00 (1.11) | 1.69 (2.05) |
| WMH % in ICV | 0.77 (0.85) | 1.33 (1.62) |
| *Ratings* |  |  |
| Fazekas score (periventricular) |  |  |
| 0 | 17 (4) | 6 (4) |
| 1 | 310 (66) | 69 (48) |
| 2 | 116 (24) | 41 (28) |
| 3 | 28 (6) | 29 (20) |
| Fazekas score (deep white matter overall) |  |  |
| 0 | 77 (16) | 22 (15) |
| 1 | 306 (65) | 79 (54) |
| 2 | 77 (16) | 21 (15) |
| 3 | 11 (2) | 23 (16) |
| PVS (basal ganglia) | Range: 0-3 | Range: 0-1* |
| 0 | 1 (0) | 87 (60) |
| 1 | 279 (60) | 57 (40) |
| 2 | 173 (37) | - |
| 3 | 14 (3) | - |
| Microbleeds |  |  |
| 0 | 420 (90) | 122 (85) |
| 1 | 47 (10) | 22 (15) |
| SVD score |  |  |
| 0 | 220 (47) | 69 (48) |
| 1 | 164 (35) | 27 (19) |
| 2 | 68 (15) | 29 (20) |
| 3 | 15 (3) | 14 (10) |
| 4 | 0 | 5 (3) |
| *Retinal measurements, mean (SD)* |  |  |
| CRAE | 31.19 (2.19) | 30.93 (2.80) |
| CRVE | 42.12 (3.16) | 43.67 (3.59) |
| BSTDa | 2.20 (0.64) | 2.10 (0.65) |
| BSTDv | 3.94 (0.86) | 4.01 (0.90) |
| FDa | 1.59 (0.05) | 1.59 (0.05) |
| FDv | 1.57 (0.04) | 1.58 (0.04) |
| TORTa (log transformed) | -9.98 (0.96) | -10.01 (0.88) |
| TORTv (log transformed) | -9.91 (0.69) | -9.76 (0.73) |
| BCa | 2.09 (0.22) | 2.14 (0.30) |
| BCv | 1.98 (0.24) | 1.98 (0.33) |
| LDRa | 19.37 (5.04) | 21.81 (11.21) |
| LDRv | 18.91 (4.96) | 19.55 (5.36) |
| AFa | 0.92 (0.04) | 0.90 (0.05) |
| AFv | 0.90 (0.05) | 0.88 (0.06) |

*Note.* BTV=brain tissue volume; ICV=intracranial volume; PVS= perivascular spaces (in basal ganglia); \*0: none-mild; 1: moderate-severe. SVD=small vessel disease; CRAE=central retinal arteriolar equivalent; CRVE=central retinal vein equivalent; BSTDa=standard deviation of arteriolar widths in Zone B; BSTDv=standard deviation of venular widths in Zone B; FDa=arteriolar fractal dimension; FDv=venular fractal dimension; TORTa=arteriolar tortuosity; TORTv=venular tortuosity; BCa=arteriolar branching coefficient; BCv=venular branching coefficient; LDRa=arteriolar length diameter ratio; LDRv=venular length diameter ratio; AFa=arteriolar asymmetry factor; AFv=venular asymmetry factor.

Prior to structural equation modelling (SEM), initial analyses in the LBC1936 used regression with 16 retinal variables as exposures and 11 brain variables as outcomes. The results of these multiple linear regression, adjusted for age and sex, are presented in Table 3a. Three of the 176 associations survived adjustment and false discovery rate (FDR) correction. Decreased arteriolar fractal dimension (decreased branching complexity) was associated with greater WMH volume percentage in both brain tissue volume (BTV) (β = -0.203, *p* < 0.001), and intracranial volume (ICV) (β = -0.203, *p* < 0.001), and higher Fazekas scores in periventricular regions (β = -0.157, *p* < 0.001). The following nominally significant age- and sex- adjusted associations attenuated to non-significance with FDR correction: arteriolar fractal dimension with Fazekas scores in deep white matter regions; venular fractal dimension with WMH % BTV and ICV; venular tortuosity with BTV mm^3^, Fazekas score in deep white matter, perivascular spaces (PVS) score and SVD score; and arteriolar branching coefficient with deep and superficial atrophy ratings, arteriolar asymmetry factor with deep atrophy ratings, and arteriolar length-diameter ratio with ICV (mm^3^).

**Table 3a.** Age- and sex- adjusted associations between retinal and brain imaging-derived measurements in Lothian Birth Cohort 1936

|  | CRAE | CRVE | BSTDa | BSTDv | FDa | FDv | TORTa | TORTv | BCa | BCv | AFa | AFv | LDRa | LDRv | Gradient<br>a<br>1<sup>st</sup><br>Unrotated<br>PC | Gradient<br>v<br>1<sup>st</sup><br>Unrotated<br>PC |
| --- | --- | --- | --- | --- | --- | --- | --- | --- | --- | --- | --- | --- | --- | --- | --- | --- |
| | $\beta$ | $\beta$ | $\beta$ | $\beta$ | $\beta$ | $\beta$ | $\beta$ | $\beta$ | $\beta$ | $\beta$ | $\beta$ | $\beta$ | $\beta$ | $\beta$ | $\beta$ | $\beta$ |
| WMH % in BTV | -.091 | -.076 | .050 | .014 | **-.203\*\*\*** | -.140<sup>a</sup> | -.027 | .084 | .030 | -.019 | .008 | .031 | .069 | .018 | .025 | .025 |
| BTV mm<sup>3</sup> | -.053 | .043 | -.089 | -.036 | .020 | -.017 | -.035 | -.113<sup>a</sup> | .029 | -.063 | -.070 | -.042 | -.031 | -.007 | .007 | .068 |
| WMH % in ICV | -.091 | -.076 | .049 | .016 | **-.203\*\*\*** | -.143<sup>a</sup> | -.027 | .082 | .031 | -.019 | .008 | .028 | .073 | .021 | .025 | .026 |
| ICV mm<sup>3</sup> | -.065 | .071 | -.099 | -.084 | .027 | .037 | -.039 | -.083 | .008 | -.101 | -.102 | .003 | -.131<sup>a</sup> | -.082 | .007 | .075 |
| Fazekas Periventricular | -.054 | .013 | .083 | .022 | **-.157\*\*\*** | -.080 | .000 | .083 | -.034 | .043 | .000 | .017 | .008 | -.003 | .012 | -.011 |
| Fazekas Deep | -.016 | -.010 | .054 | .056 | -.118<sup>a</sup> | -.084 | .008 | .113<sup>a</sup> | -.010 | .023 | .014 | -.005 | .060 | .044 | .022 | .059 |
| Atrophy Deep | -.035 | -.007 | .024 | -.011 | -.076 | .021 | .021 | .073 | .113<sup>a</sup> | .075 | -.100<sup>a</sup> | -.032 | -.038 | -.058 | .042 | -.019 |
| Atrophy Superficial | -.017 | -.007 | .044 | -.041 | -.071 | .026 | .011 | .046 | .112<sup>a</sup> | .076 | -.082 | -.001 | -.062 | -.072 | -.024 | -.014 |
| PVS | -.043 | .028 | -.014 | -.044 | .041 | .080 | .010 | .146<sup>a</sup> | -.005 | -.016 | .047 | .038 | -.014 | .041 | .002 | -.029 |
| Microbleeds | -.020 | .003 | -.052 | .000 | .000 | .016 | .028 | -.018 | -.062 | -.024 | .025 | .032 | .019 | -.067 | -.039 | -.048 |
| SVD score | -.059 | .009 | -.005 | -.009 | -.051 | .025 | .007 | .141<sup>a</sup> | -.029 | .018 | .040 | .024 | .033 | .001 | .037 | -.005 |

*Note.* Model adjusted for age and sex. WMH=white matter hyperintensities; BTV=brain tissue volume; ICV=intracranial volume; PVS= basal ganglia perivascular spaces; SVD=small vessel disease; CRAE=central retinal artery equivalent; CRVE=central vein retinal equivalent; BSTDa=standard deviation of arteriolar widths in Zone B; BSTDv=standard deviation of venular widths in Zone B; FDa= fractal dimension arteriolar; FDv=fractal dimension venular; TORTa= arteriolar tortuosity; TORTv=venular tortuosity; BCa=arteriolar branching coefficient; BCv=venular branching coefficient; AFa=arteriolar asymmetry factor; AFv=venular asymmetry factor; LDRa=arteriolar length diameter ratio; LRDv= venular length diameter ratio; Gradient a= arteriolar width gradient; Gradient v= venular width gradient; PC=principal component; Bolded values are statistically significant. WML % volumes and tortuosity log transformed. $\beta$ , Standardised regression coefficients.
\* = $p < .05$ ; \*\* = $p < .01$ ; \*\*\* = $p < .001$ ; all p-values corrected for False Discovery Rate. <sup>a</sup> = value was statistically significant at $p < .05$ before FDR correction

The above analyses were re-run after excluding LBC1936 participants with history of stroke (n=84) (Supplementary Table S1). Two additional associations survived adjustment and FDR correction in this sensitivity analysis; decreased venular fractal dimension was associated with increased WMH % in BTV (β = -0.175, *p* = 0.03) and ICV (β = -0.178, *p* = 0.03). The three other previously-significant associations with arteriolar fractal dimension remained significant.

Measurement models in SEM to derive the latent traits of VRF, ‘WMH load’, ‘SVD burden’ and retinal ‘calibre-complexity’ in the LBC1936 sample (Supplementary Fig. S1-S3) indicate that almost all measured variables contributed significantly to their hypothesised latent variable (with parameter weights with *p* < 0.05). The exceptions were: smoking (*p* = 0.34), as found in the VRF latent variable derived previously in the LBC1936^16^; and microbleeds in the SVD latent trait (*p* = 0.29). Nonetheless, these variables were included in the VRF and SVD latent traits. All models had acceptable fit to the data (Supplementary Fig. S1-S3).

SEMs used to test hypothesized relationships among latent variables of ‘calibre-complexity’ and ‘WMH load’ and ‘SVD burden’ in LBC1936 are shown in Figure 1. In addition to the contributions of age, sex and VRFs, ‘calibre-complexity’ was significantly associated with the latent traits of WMH load (standardised parameter weight -0.181, *p* < 0.001) (Figure 1A) and SVD burden (standardised parameter weight -0.141, *p =* 0.021) (Figure 1B). This retinal construct of vessel calibre and branching complexity accounted for approximately 3.3% and 2.0% of the variance in WMH load and SVD burden, respectively. However, the association between ‘calibre-complexity’ and SVD burden attenuated to non-significance once corrected for multiple comparisons (*p* = 0.054).

**Figure 1.**
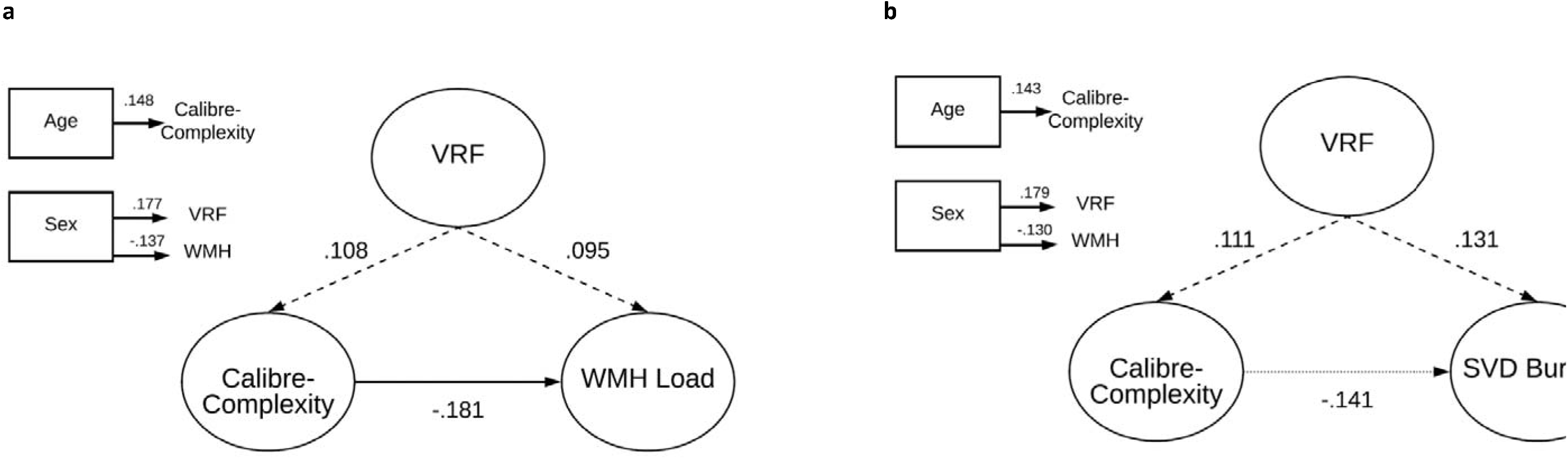
Structural models for retinal latent traits predicting WMH/SVD burden in the LBC1936 (a) Model of retinal calibre-complexity predicting WMH Load; (b) Model of retinal calibre-complexity predicting SVD burden; Standardised regression coefficients (parameter weights) are shown adjacent to each path. P-values corrected for False Discovery Rate. The rectangles to the left are the covariates in the model. Each of the covariates was examined for their contribution to VRF, ‘Calibre-complexity’ and WMH Load/SVD Burden, and only significant (*p* < .05) contributions are shown. VRF = vascular risk factor; WMH = white matter hyperintensities; SVD = small vessel disease. Dashed lines represent non-significant associations; dotted line represents nominally significant association.

SEMs used to test hypothesized relationships among individual retinal variables (identified from age- and sex- adjusted regression models, see Table 3), and latent constructs of WMH load and SVD burden in LBC1936 are shown in Figures 2. In addition to the contributions of age, sex and VRFs, reduced arteriolar fractal dimension was significantly associated with the latent traits of WMH load (standardised parameter weight -0.211, *p* < 0.001) (Figure 2A) and SVD burden (standardised parameter weight -0.168, p = 0.002) (Figure 2B), accounting for approximately 4.4% and 2.8% of the variance in WMH and SVD respectively. These associations survived FDR correction. The VRF latent trait was not significantly associated with fractal dimension or either SVD or WMH latent trait (Figure 2).

**Figure 2.**
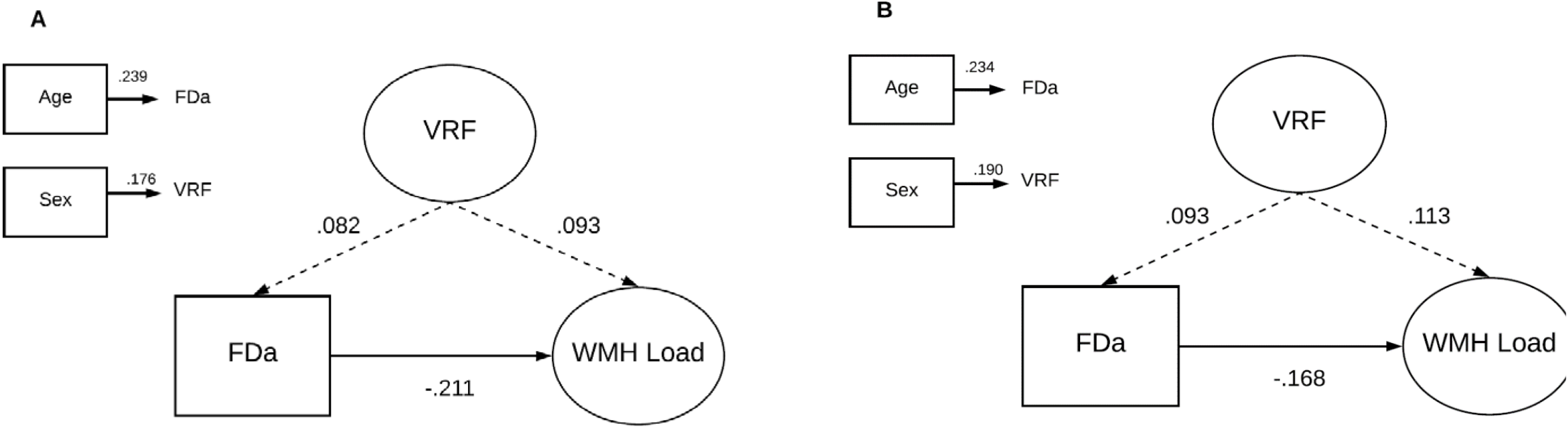
Structural models for retinal fractal dimension measurements predicting WMH/SVD burden in the LBC1936 (A) Model of FDa predicting WMH Load; (B) Model of FDa predicting SVD burden; Standardised regression coefficients (parameter weights) are shown adjacent to each path. P-values corrected for False Discovery Rate. The rectangles to the left are the covariates in the model. Each of the covariates was examined for their contribution to VRF, FDa and WMH Load/SVD Burden, and only significant (*p* < .05) contributions are shown. FDa = arteriolar fractal dimension; VRF = vascular risk factor; WMH = white matter hyperintensities; SVD = small vessel disease. Dashed lines represent non-significant associations.

We tested additional SEMs—to test for co-association of retinal and SVD indicators with VRFs—of the three relationships between retinal variables and individually-measured brain imaging variables that had surviving age-, sex-, and FDR adjustment (Table 3a). The results are shown in Supplementary Fig. S13. All retinal-brain associations remained significant and survived FDR correction.

### MSS

Among those participants of the MSS cohort with retinal images of both eyes available for analysis (n=155; 98, 68% male), of mean age 66.9 years (SD 11.4), 87 (60%) had hypertension, 19 (13%) had diabetes (Table 1). Seventy (48%) had moderate or severe periventricular WMH scores, and 44 (30%) with moderate or severe deep WMH scores (Table 2).

In age- and sex-adjusted linear regression models, only one association remained significant following correction for FDR; reduced arteriolar fractal dimension was significantly associated with higher WMH scores in deep white matter regions (β = -0.327, *p* < 0.001). The following 12 nominally significant age- and sex- adjusted associations attenuated to non-significance after applying FDR correction: central retinal artery equivalent (CRAE) with WMH % BTV and ICV, variation in arteriolar widths with WMH % BTV and ICV; variation in venular widths with WMH % in BTV and ICV and Fazekas scores in deep white matter; arteriolar fractal dimension with WMH % in BTV, ICV, Fazekas scores in periventricular region, and SVD score; and venular branching coefficient and Fazekas score in deep white matter region (Table 3b).

**Table 3b.**
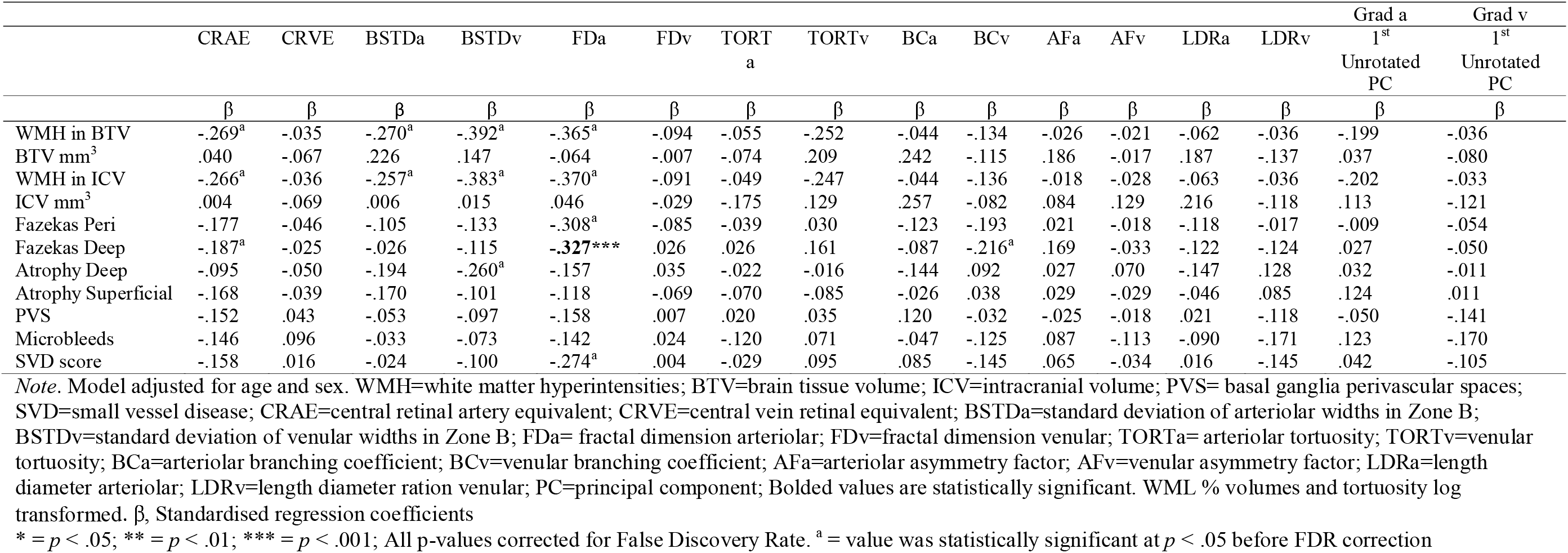
Age- and sex- adjusted associations between retinal and brain imaging-derived measurements in MSS

|  | CRAE | CRVE | BSTDa | BSTDv | FDa | FDv | TORT<sub>a</sub> | TORT<sub>v</sub> | BCa | BCv | AFa | AFv | LDRa | LDRv | Grad a<br>1<sup>st</sup><br>Unrotated<br>PC | Grad v<br>1<sup>st</sup><br>Unrotated<br>PC |
| --- | --- | --- | --- | --- | --- | --- | --- | --- | --- | --- | --- | --- | --- | --- | --- | --- |
|  | β | β | β | β | β | β | β | β | β | β | β | β | β | β | β | β |
| WMH in BTV | -.269<sup>a</sup> | -.035 | -.270<sup>a</sup> | -.392<sup>a</sup> | -.365<sup>a</sup> | -.094 | -.055 | -.252 | -.044 | -.134 | -.026 | -.021 | -.062 | -.036 | -.199 | -.036 |
| BTV mm<sup>3</sup> | .040 | -.067 | .226 | .147 | -.064 | -.007 | -.074 | .209 | .242 | -.115 | .186 | -.017 | .187 | -.137 | .037 | -.080 |
| WMH in ICV | -.266<sup>a</sup> | -.036 | -.257<sup>a</sup> | -.383<sup>a</sup> | -.370<sup>a</sup> | -.091 | -.049 | -.247 | -.044 | -.136 | -.018 | -.028 | -.063 | -.036 | -.202 | -.033 |
| ICV mm<sup>3</sup> | .004 | -.069 | .006 | .015 | .046 | -.029 | -.175 | .129 | .257 | -.082 | .084 | .129 | .216 | -.118 | .113 | -.121 |
| Fazekas Peri | -.177 | -.046 | -.105 | -.133 | -.308<sup>a</sup> | -.085 | -.039 | .030 | -.123 | -.193 | .021 | -.018 | -.118 | -.017 | -.009 | -.054 |
| Fazekas Deep | -.187<sup>a</sup> | -.025 | -.026 | -.115 | **-.327\*\*\*** | .026 | .026 | .161 | -.087 | -.216<sup>a</sup> | .169 | -.033 | -.122 | -.124 | .027 | -.050 |
| Atrophy Deep | -.095 | -.050 | -.194 | -.260<sup>a</sup> | -.157 | .035 | -.022 | -.016 | -.144 | .092 | .027 | .070 | -.147 | .128 | .032 | -.011 |
| Atrophy Superficial | -.168 | -.039 | -.170 | -.101 | -.118 | -.069 | -.070 | -.085 | -.026 | .038 | .029 | -.029 | -.046 | .085 | .124 | .011 |
| PVS | -.152 | .043 | -.053 | -.097 | -.158 | .007 | .020 | .035 | .120 | -.032 | -.025 | -.018 | .021 | -.118 | -.050 | -.141 |
| Microbleeds | -.146 | .096 | -.033 | -.073 | -.142 | .024 | -.120 | .071 | -.047 | -.125 | .087 | -.113 | -.090 | -.171 | .123 | -.170 |
| SVD score | -.158 | .016 | -.024 | -.100 | -.274<sup>a</sup> | .004 | -.029 | .095 | .085 | -.145 | .065 | -.034 | .016 | -.145 | .042 | -.105 |

*Note.* Model adjusted for age and sex. WMH=white matter hyperintensities; BTV=brain tissue volume; ICV=intracranial volume; PVS= basal ganglia perivascular spaces; SVD=small vessel disease; CRAE=central retinal artery equivalent; CRVE=central vein retinal equivalent; BSTDa=standard deviation of arteriolar widths in Zone B; BSTDv=standard deviation of venular widths in Zone B; FDa= fractal dimension arteriolar; FDv=fractal dimension venular; TORTa= arteriolar tortuosity; TORTv=venular tortuosity; BCa=arteriolar branching coefficient; BCv=venular branching coefficient; AFa=arteriolar asymmetry factor; AFv=venular asymmetry factor; LDRa=length diameter arteriolar; LDRv=length diameter ration venular; PC=principal component; Bolded values are statistically significant. WML % volumes and tortuosity log transformed. β, Standardised regression coefficients
\* = $p < .05$ ; \*\* = $p < .01$ ; \*\*\* = $p < .001$ ; All p-values corrected for False Discovery Rate. <sup>a</sup> = value was statistically significant at $p < .05$ before FDR correction

We conducted further multivariable ordinal regression analysis on the surviving association between arteriolar fractal dimension and deep white matter Fazekas score, introducing the VRF component as an additional covariate to age and sex (Table 4). Lower arteriolar fractal dimension remained significantly associated with deep WMH Fazekas scores, with OR (95% CI) of 0.53 (0.32-0.87).

**Table 4.** Multivariate regression analysis of brain imaging-derived measurements of SVD in the Mild Stroke Study. All variables in column 1 were entered simultaneously as independent variables in the models

|  | *Fazekas Deep*<sup>†</sup> |  |
| --- | --- | --- |
|  | OR (95% CI) | *p* |
| Male | 1.23 (0.49; 3.42) | 0.596 |
| Age | 2.30 (1.45; 3.65) | <.000 |
| VRF | 0.85 (0.54; 1.33) | 0.474 |
| FDa | 0.53 (0.32; 0.87) | 0.012 |

*Note.* Multivariate analysis with correction for all variables in the table. OR=odds ratio; CI=confidence interval; FDa=arteriolar fractal dimension; SVD=small vessel disease. <sup>†</sup> Ordinal regression with Fazekas score (0-3) as dependent variable. Continuous predictors were converted to z-scores.

## Discussion

To our knowledge, the current study is the first that uses SEM to investigate the relationship between retinal microvascular and brain imaging markers of SVD. Whereas our retinal data reduction approach found most retinal measurements to be largely independent, i.e. not indicators of the same latent trait, we did find four retinal variables with considerable shared variance reflecting aspects of geometry, namely vessel width and complexity. This combined vessel calibre-complexity latent trait was related to SVD burden and WMH load in the LBC1936. Examining retinal variables individually in the LBC1936, we found participants with decreased arteriolar fractal dimension, suggesting of a loss of branching complexity, were more likely to have greater cerebral SVD pathology, independent of VRF. Sparser arteriolar microvasculature was also associated with more WMH. All effect sizes were small. The relatively larger effect sizes of the results using individual retinal variables suggest that the relationship between ‘calibre-complexity’ and SVD burden is largely driven by the effect of reduced arteriolar fractal dimension. Findings were generally consistent in the MSS participants despite the wider age range and increased burden of vascular disease; MSS participants with sparser arteriolar fractal dimension were more likely to have more WMH in deep white matter. Our results are in line with those of Wardlaw et al. (2014) in reporting that VRFs only account for a small proportion of the variance in WMH^16^. Our findings add further weight to the concept that easy to measure, common VRFs explain only a small proportion of both variance in retinal vascular and brain SVD features, leaving a large proportion unexplained.

These findings add further support to the hypothesis that retinal fractal dimension, as a global summary measure of the state or health of the retinal vasculature, might have a significant association with the changes taking place in cerebral small vessels. A loss of branching complexity in regulatory systems of other organs of the body is thought to occur as they become less complex with age^17^. In line with our results, Hilal et al. (2014) found those with reduced arteriolar fractal dimension were more likely to have cerebral microbleeds^11^. Reduced fractal dimension has also been found in Alzheimer’s disease^18-20^. Decreased branching complexity of the arteriolar and venular networks combined in lacunar stroke, a manifestation of SVD, has previously been reported in the MSS cohort^6^.

A large number of association tests were conducted, and several significant associations became non-significant once corrected for multiple testing. Whereas the amount of associations conducted increases the likelihood of type-1 errors, it is also possible that our robust correction for multiple testing is rejecting modest but true associations; larger sample sizes are needed to confirm them^21^. We interpret all unadjusted significant associations cautiously as they may reflect spurious associations. However, they could provide directions for future research. Without adjustment, some additional retinal measurements, including venular tortuosity and arteriolar measures of branching geometry, appeared relevant for some markers of SVD.

A limitation of the study is its cross-sectional design. Longitudinal studies examining retinal changes and progression of SVD and WMH would be valuable. The smaller sample size of the MSS cohort did not allow for consistent statistical methods across cohorts. Furthermore, despite the use of a validated semi-automated software application; VAMPIRE (Vessel Assessment and Measurement Platform for Images of the Retina, version 3.1, Universities of Edinburgh and Dundee), a degree of subjective human interaction introduces potential for intra-grader variability. Sources of variability of retinal measurements from fundus images have been discussed, among others, in Trucco et al. (2013)^22^. We also acknowledge our retinal variables data reduction approach represents a first step towards a composite measure of abnormality and subsequent choice of variables is not definitive. Other studies may reach different conclusions based on their data. The participants’ age may have reduced the strength of the associations between VRF, WMH and retinal measurements. The relationship between WMH and hypertension appears stronger in middle age^23^. A systematic review of associations between retinal microvascular changes, dementia and brain imaging features reported stronger findings in middle-aged populations^24^, in line with the age-related attenuation of associations between VRFs, hypertension and cerebrovascular disease and retinopathy^25, 26^.

The strengths of the current study include the use of two distinct, yet complementary, well-characterised cohorts. This provides a more comprehensive account of the relationship between the retinal and cerebral microvasculature in different populations, representing different points on the SVD continuum. Furthermore, the LBC1936 as a narrow-age cohort, reduces the troublesome problem where chronological age confounds the influence of other variables. All LBC1936 and MSS participants were imaged using the same retinal camera and MRI scanner, at the same time period, according to the same brain and retinal image analysis methods. This included the use of standardised retinal and cerebral examinations and the use of both quantitative and qualitative WMH measures with the careful exclusion of infarcts from WMH volume. Additional strengths include the use of robust statistics. The sample size in the LBC1936 allowed us to account for measurement error by using SEM to derive error-free latent contracts for VRF, WMH and SVD.

In summary, we provide some evidence to support concomitant retinal and cerebral microvasculature changes, with a small but significant association between reduced retinal arteriolar branching complexity and SVD pathology, known to represent a significant risk factor for dementia and stroke^27^. That sparser vasculature is associated with WMH and SVD burden, suggests that the retinal microvasculature can exhibit corresponding microvascular signs of altered vascular competence leading to tissue damage in the brain. These data extend the research potential of retinal imaging beyond the eye in studying neurovascular disease^1^ and paves the way for further studies into mechanism of dementia and stroke and possibly to refine the prognostic potential of detailed retinal microvascular analysis, although we again note that the associations are small. The added value of embedding retinal image analytics in cross-linked health datasets to study SVD is yet to be determined; this requires intra-individual change detection in stratified prospective cohorts to be tracked. As a cross-sectional study, we can only determine associations not causation, nor the progression of cerebral SVD. Despite our efforts in judging the statistical significance of the results with FDR correction, the associations found in this study need future replication in independent samples. Careful validation of our findings is required for better characterizing the retinal changes accompanying SVD, and in considering whether retinal measures have a place in clinical work such as in screening or as outcomes in trials. Further studies are required to elucidate whether changes to retinal properties precede pathology or a response to long-term exposure to risk factors.

## Methods

### Participants

The LBC1936^28^ comprised community-dwelling, mostly healthy older adults, of mean age about 70 years when first recruited in older age. All were born in 1936. The data analysed for the present study, including digital retinal photographs and structural brain imaging, were obtained at a second wave of testing when the participants were approximately 73 years old (N=866). The recruitment and testing of the LBC1936 has been described in detail previously^28-30^.

The Mild Stroke Study (MSS)^4^ is a prospective study of patients with recent (within 3 months) clinical lacunar or mild cortical ischaemic stroke. All patients were assessed by an experienced stroke physician. The recruitment, testing and imaging of these patients has been described previously^4,10^.

Both studies were approved by Lothian Research Ethics (LBC1936: REC 07/MRE00/58; MSS: 2002/8/64). The LBC1936 study was also approved by the Scottish Multicentre (MREC/01/0/58) Research Ethics. Written informed consent for participation in both studies was obtained from all participants. The research was carried out in compliance with the Helsinki Declaration.

### Measures

#### Retinal image acquisition and analysis

In both groups, digital retinal fundus images were captured using the same non-mydriatic camera at 45° field of view (CRDGi; Canon USA, Lake Success, New York, USA). 814 LBC1936 (from Wave 2 of testing) and 190 MSS participants provided retinal images of both eyes. Images were centred approximately on the optic disc. For the present analysis, the retinal images were reanalysed for retinal vascular characteristics using the same semi-automated software package, VAMPIRE by an experienced operator. VAMPIRE image processing and analysis has been described in detail previously^31-33^. Briefly, the boundaries of the optic disc (OD) and position of the fovea in a retinal image are first detected and the conventional set of OD-centred circular measurement zones established. Zone B is a ring 0.5 to 1 OD diameters away from the centre, and Zone C is a ring extending from OD boundary to 2 OD diameters away. Next, the software detects the retinal blood vessels present in the image and classifies them as arterioles or venules. The observer, when necessary followed a standardised measurement protocol to preform manual interventions to correct computer-generated labelling of image features, blind to all prior retinal analysis, brain and VRF data. There were complete retinal measurements from both eyes for 603 LBC1936 and 155 MSS participants. Rejections were due to poor quality images, eyelashes causing streaks across the photograph, out-of-focus images, and overexposure (in either eye); these occurred in approximately 16% of LBC1936 images, and 8% of MSS images, with an additional 4% of MSS images excluded due to considerable differences in image resolution (arising from deviations from standard operation of the system when imaging).

Sixteen retinal vascular parameters were measured from each image in both cohorts: measures of vessel calibre—central retinal artery equivalent (CRAE), central retinal vein equivalent (CRVE), and the variation in calibre—the standard deviation of arteriolar and venular widths (BSTDa, BSTDv); the gradient of the width of the main arteriolar and venular vessel paths (GRADa, GRADv); measures of branching complexity—arteriolar and venular fractal dimension (FDa, FDv); measures of vessel tortuosity—arteriolar and venular tortuosity (TORTa, TORTv); and measures of arteriolar and venular branching geometry— branching coefficient (BCa, BCv), length-diameter ratio (LDRa, LDRv) and asymmetry factor (AFa, AFv). A lowercase ‘a’ or ‘v’ following the variable name indicates a measurement of arteriolar or venular vessels respectively. See supplementary material for details on all retinal measurements and how retinal variables were selected for analysis. To reduce the number of variables, reduce multicollinearity and increase reliability, the above-mentioned measurements from both eyes of each participant were averaged to provide a mean measurement for all variables.

#### MRI brain image acquisition and processing

LBC1936 and MSS participants (at time of presentation) underwent brain MRI on the same 1.5-Tesla GE Signa Horizon HDx clinical scanner (General Electric, Milwaukee, WI) with T1-, T2- and T2*- weighted and fluid-attenuated inversion recovery (FLAIR) axial whole-brain imaging. Full details of the brain imaging scanning protocol for the LBC1936 and MSS have been described previously^10,30^. All analyses were performed blinded to all other data. The SVD lesions in both studies were assessed qualitatively and quantitatively using validated methods according to a precursor to the STRIVE criteria^34^. WMH were visually scored using FLAIR-, T1 and T2-weighted sequences on the Fazekas score^35^ in both the deep (0-3) and periventricular (0-3) white matter. Appropriate sequences were also rated for the presence of microbleeds (location and number), lacunes (location and number), and perivascular spaces (in basal ganglia and centrum semiovale, 0-4 point score each) according to an established rating protocol^36^. Brain atrophy was coded using a validated template^37^, with superficial and deep atrophy coded separately.

We combined the visual lesion scores into an ordinal ‘total SVD score’ of 0-4, described previously^38^. Briefly, a scale point was awarded for the presence of (early) confluent deep (2- 3) WMH and/or irregular periventricular WMH extended into the deep white matter (3); one or more lacunes; one or more microbleeds; and moderate to severe grading (2-4) of basal ganglia perivascular spaces. These showed face-validity both as an ordinal score and as a latent variable in previous analyses both in the present cohorts and in other studies^38,39^. All rating was performed by a consultant neuroradiologist trained and experienced in SVD features and use of the visual ratings. Quality control of images has been described previously^16,39^.

Quantitative measures of WMH, brain and intracranial volume were obtained using T2^*^- weighted and FLAIR sequences with a validated semi-automated multispectral image processing tool, MCMxxxVI^40^. This tool was used to measure intracranial volume (ICV, soft tissue structures within the cranial cavity including brain, cerebrospinal fluid, dural and venous sinuses), brain tissue volume (BTV, intracranial volume excluding the ventricular cerebrospinal fluid) and WMH. The structure volumes were measured as absolute values in cubic millimetres (BTV mm^3^, ICV mm^3^). Quantitative measures of WMH were expressed as percentage of WMH volume in ICV (WMH % ICV) and percentage of WMH volume in BTV (WMH % BTV).

#### Covariates

Age and sex were included as covariates in both the LBC1936 and MSS samples. Measures of vascular risk were included as covariates in both samples. VRFs were assessed in the LBC1936 subjects at age ~73 years, at the same session as the retinal photography, and a mean (SD) of 9 (5) weeks prior to brain imaging; they were assessed on presentation in the MSS, at the same time as brain imaging, and approximately four weeks prior to retinal photography. A combination of medical history variables (medically diagnosed hypertension, diabetes, smoking, and hypercholesterolemia), and measured variables (blood pressure [BP], haemoglobin A1c, and plasma cholesterol) were used. The average of three sitting BP measurements were used to derive mean systolic and mean diastolic BP variables in LBC1936 and one BP reading was used for MSS subjects. The above measures were recorded for MSS subjects with the exception of haemoglobin A1c. All measures were performed blinded to all other data. Variables were selected based on a set of measures of vascular risk that we had identified contributed to vascular risk of WMH in previous LBC1936 and MSS analysis^16^.

### Statistical analysis

Age- and sex-adjusted linear regression was used to analyse the association between the 16 retinal vascular characteristics and the structural brain imaging-derived measurements in both cohorts. To minimise the potential for type I errors, p values were adjusted according to the false discovery rate (FDR) method^41^. LBC1936 participants with a history of stroke (n=84, 14%; based on medical history and/or brain imaging appearances) were removed in a sensitivity analysis. Due to the small size and insufficient stroke classification this group could not be divided into stroke subtypes. VRFs were tested as possible explanatory variables for any significant associations between retinal and brain imaging variables, since both retinal vascular abnormalities and SVD features are known to be associated with common VRFs such as hypertension, smoking, diabetes, etc.; this was examined using SEM in LBC1936, and multivariable regression models in the MSS cohort (which we judged to be too small for SEM). See Penke and Deary (2010) for an accessible description of SEM as applied in neuroscience^42^.

The basic questions in the analyses were whether retinal vessel measures were associated with brain imaging measures, and whether these associations were co-associated with VRFs. The LBC1936 is both large in size and has multiple measures of brain white matter health and VRFs. Therefore, in testing the questions above, we were able to form multi-variable ‘latent traits’ (unobservable constructs underlying a combination of correlated individual measured variables) for retinal features, white matter health, and vascular risk. Results from the regression analyses in the LBC1936 were used to motivate the hypothesized relationships subsequently to be tested by SEM.

We showed previously that VRFs, WMH measures and SVD features formed latent variables in the LBC1936^16, 39,43^. Therefore, we used the same measurement models to derive the latent variables. Vascular risk was measured as a single latent factor from eight variables; hypertension, diabetes, hypercholesterolemia, smoking, (treated) systolic and diastolic BP, haemoglobin A1c, and plasma cholesterol, as previously^16^. The volume of WMH as a percentage of ICV, and Fazekas ratings in periventricular and deep white matter were used to derive a latent variable of ‘WMH load’ as previously^43^. ‘SVD burden’ was measured using a single latent factor with five indicators, namely, Fazekas ratings for both periventricular and deep regions, lacunes, microbleeds, and basal ganglia perivascular spaces, as previously^39^. This was undertaken to test whether including three additional imaging markers of SVD might increase the ability to find significant associations. A single latent ‘calibre-complexity’ factor was derived from four retinal indicators; two measures of vessel width (CRAE, CRVE), and two measures of branching complexity; arteriolar and venular fractal dimension. The derivation of this latent variable is described fully in the Supplementary Material. All models were estimated using R’s lavaan SEM package, version 0.5-22^44^.

Models were estimated using the robust (mean and variance adjusted) weighted least squares (WLSMV) estimator. WLSMV is robust to non-normality and is appropriate for model estimation with categorical data. Standardised regression coefficients (parameter weights, comparable to standardised partial beta weights) were computed for each path in the models. Model fit was assessed using cut-off points of > 0.06 for the root mean square error of approximation (RMSEA), and ≥ 0.90 for the comparative fit index (CFI) and Tucker-Lewis index (TLI). Measurement models for latent traits are shown in Supplementary Fig. S1-S3.

We tested the same two questions in the MSS as above for the LBC1936. However, due to the smaller sample size, latent variables were not formed in MSS and we did not use SEM to test hypotheses. Instead, multivariable regression models were applied in the MSS to test for associations between retinal and brain imaging-derived measurements, and the controls were applied for age, sex, and VRFs. To reduce the number of vascular risk parameters and the likelihood of type I errors, principal components analysis (PCA) was applied to the eight measured VRF variables in MSS. The first unrotated principal component accounted for a substantial percentage of the overall variance in VRF variables (26%), with loadings ranging between 0.18 and 0.77, and was used to generate a general VRF score. To validate the use of a principal component score, factor scores for VRFs were derived using the same PCA method in the LBC1936 sample. The correlation between the VRF principal component score from PCA analysis and the VRF latent trait obtained using SEM in the LBC1936 was very strong (*r* = 0.89). Multivariable ordinal regression analysis was used for WMH and SVD scores in MSS. Results are presented as odds ratio (OR) with 95% confidence interval (CI). Predictors were converted to z-scores, such that the resulting ORs reflect the odds of having higher pathology ratings for each standard unit increase in the predictor variable. Regression analyses were performed with SPSS statistics version 22 (IBM Corp., Armonk, NY).

## Acknowledgements

The authors thank the Lothian Birth Cohort 1936 members and participants of the Mild Stroke Study who took part in this study; the radiographers at the Brain Research Imaging Centre; the nurses of the Wellcome Trust Clinical Research Facility and the members of the LBC1936 research team who collected and collated the LBC1936 data we analysed. The Lothian Birth Cohort 1936 is supported by Age UK (Disconnected Mind project) and the Medical Research Council (MR/M01311/1). Support from NHS Lothian R&D, and Edinburgh Imaging and Edinburgh Clinical Research Facility at the University of Edinburgh is gratefully acknowledged. Retinal photographs were taken in the Wellcome Trust Clinical Research Facility, Western General Hospital, Edinburgh. We thank Stuart J. Ritchie for statistical advice and guidance. The work was undertaken in The University of Edinburgh Centre for Cognitive Ageing and Cognitive Epidemiology, which is supported by funding from BBSRC and the Medical Research Council (MRC) as part of the cross council Lifelong Health and Wellbeing Initiative (MR/K026992/1). The work was partially supported by EPSRC project EP/M005976/1 “Multimodal retinal biomarkers for vascular dementia”.

## Data availability

The datasets generated during and/or analysed during the current study are available from the corresponding author on reasonable request.

## Author contributions

The author contributions are as follows for each of the categories listed. Study conception and design, acquisition/analysis/interpretation of data: SM, IJD, JMW, FND, TJM, ET. Image processing: SM, MVH, XW, MEB. Drafting the article or revising it critically: SM, LB, FND, JS, MA, MVH, XW, TJM, ASFD, BD, JMS, MEB, ET, IJD, JMW

